# BATF relieves hepatic steatosis by inhibiting PD1 and promoting energy metabolism

**DOI:** 10.1101/2023.04.18.537352

**Authors:** Zhiwang Zhang, Qichao Liao, Tingli Pan, Lin Yu, Zupeng Luo, Songtao Su, Shi Liu, Menglong Hou, Yixing Li, Turtushikh Damba, Yunxiao Liang, Lei Zhou

## Abstract

The rising prevalence of nonalcoholic fatty liver disease (NAFLD) has become a global health threat that needs to be addressed urgently. Basic leucine zipper ATF-like transcription factor (BATF) is commonly thought to be involved in immunity, but its effect on lipid metabolism is not clear. Here, we investigated the function of BATF in hepatic lipid metabolism. BATF alleviated high-fat diet (HFD)-induced hepatic steatosis and inhibited elevated programmed cell death protein (PD)1 expression induced by HFD. A mechanistic study confirmed that BATF regulated fat accumulation by inhibiting PD1 expression and promoting energy metabolism. PD1 antibodies alleviated hepatic lipid deposition. In conclusion, we identified the regulatory role of BATF in hepatic lipid metabolism and that PD1 is a target for alleviation of NAFLD. This study provides new insights into the relationship between BATF, PD1 and NAFLD.

## Introduction

Nonalcoholic fatty liver disease (NAFLD) has become a prevalent chronic liver disease that threatens human health globally. Studies have shown that the global prevalence of NAFLD is about 25% and this trend is still rising^1^. The clinical diagnostic criterion for NAFLD is that >5% of the liver has steatosis^2^. Further deterioration of NAFLD leads to nonalcoholic steatohepatitis (NASH)^3^ and liver fibrosis, markedly increased risk of adverse outcomes including overall mortality, and liver-specific morbidity and mortality, respectively^4^.

Transcription factors (TFs) are a class of DNA-binding proteins whose gene regulation ability is critical to the molecular state of cells^5^. Multiple studies have shown that TFs are closely related to NAFLD. The currently discovered TFs related to liver lipid metabolism include peroxisome proliferator-activated receptors (PPARs)^6^, liver X receptors (LXRs)^7^ and sterol-regulatory element binding proteins (SREBPs)^8^. These TFs are essential for maintaining the body’s lipid metabolism balance through the regulation of lipid metabolism genes^9^. Drugs targeting TFs have been developed, and are widely used in the prevention and treatment of lipid metabolic disorders such as obesity, hyperlipidemia, diabetes, etc., and show excellent clinical effects and potential applications^10–11^.

Basic leucine zipper ATF-like transcription factor (BATF), BATF2 and BATF3 belong to the activator protein (AP)-1 family^12^. Initially, BATF family members were only considered to be inhibitors of AP-1-driven transcription, but recent studies have found that these TFs have unique transcriptional activities in dendritic cells, B cells and T cells^13^. This indicates that their functions may not be fully understood. Currently, BATF has not been found to be directly related to lipid metabolism. The purpose of this study was to explore this relationship. Our study provides a new perspective for understanding the occurrence of NAFLD.

## Results

### BATF reduces high-fat-diet-induced lipid accumulation in hepatocytes

High-fat diet (HFD) promotes hepatic lipid accumulation and leads to dysregulation of lipid metabolism. We detected hepatic BATF expression in mice fed an HFD for 14 weeks to investigate the role of BATF in hepatic lipid metabolism. HFD significantly increased liver lipid deposition and BATF levels in mice (Fig 1A, B). HepG2 cells were treated with oleic acid/palmitic acid (OA/PA) to mimic HFD. It promoted triglyceride (TG) accumulation (Fig S1A) and increased BATF expression (Fig 1C). Our analysis of data from patients with NAFLD showed that BATF levels increased with increasing NAFLD score (Fig 1D). We inhibited BATF in HepG2 cells (Fig S1B) and found no effect on cellular TG accumulation (Fig 1E). This indicated that inhibition of BATF could not regulate lipid deposition in hepatocytes (Fig 1F). Therefore, the role of BATF overexpression was studied. Oil red O staining on lipid droplets and TG detection showed that BATF alleviated OA/PA-induced HepG2 cellular fat deposition (Fig 1G, H). BATF also reduced TG accumulation in L02 human hepatocytes (Fig 1I, J). To confirm the results, we isolated mouse primary hepatocytes and infected them with a virus expressing BATF. The results also showed that BATF alleviated accumulation of lipid droplets and TG (Fig 1K, L). In AML12 cells, we found consistent results (Fig S1C, D). These results confirm that BATF can alleviate lipid deposition in hepatocytes.

**Figure 1.**
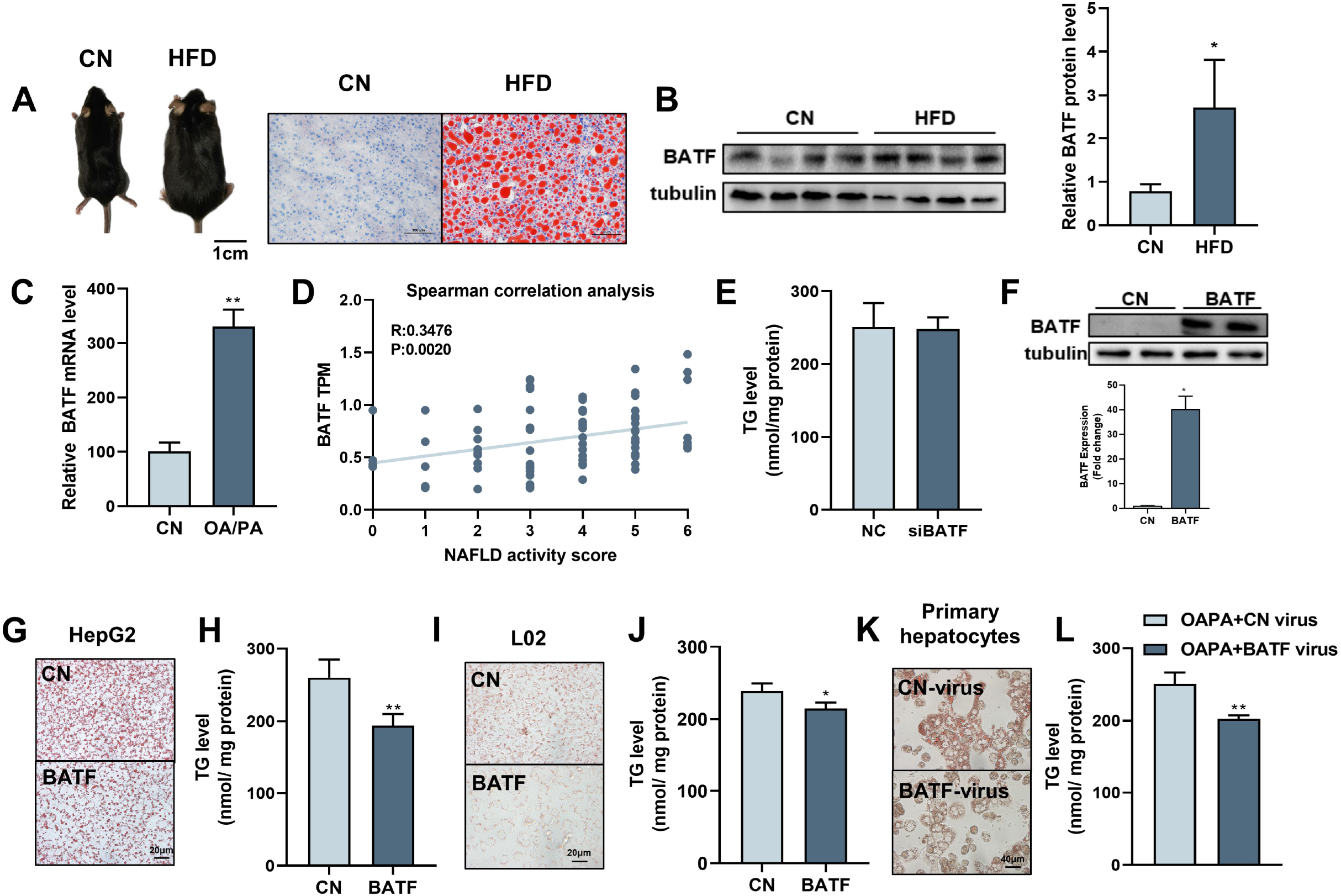
Effects of BATF on lipid deposition in hepatocytes under high-fat diet. (A) The mice and liver Oil red O staining in normal diet group (CN) and high-fat diet group (HFD). Bar, 1 cm (left panel) and 100 μm (right panel). (B) The protein expression of BATF in liver tissues, (n = 4). (C) The mRNA expression of BATF in liver tissues (n = 5). (D) Spearman correlation Analysis between TPM of BATF and NAFLD Patients with Different Degrees, (n = 4-18). (E) Triglyceride content (n = 5). (F) Detection of BATF overexpression in HepG2. (G) Oil red O staining of HepG2 cells with OA/PA when BATF was overexpressed and (H) triglyceride content, (n = 4). (I) Oil red O staining of L02 cells with OA/PA when BATF was overexpressed and (J) triglyceride content, (n = 3). (K) Oil red O staining of primary hepatocytes with OA/PA when BATF was overexpressed and (L) triglyceride content, (n = 3). The data are expressed as mean ± SD. *P < 0.05 **P < 0.01.

### Increased hepatic BATF alleviates HFD-induced hepatic steatosis in mice

We constructed an AAV8 virus overexpressing BATF to infect mouse liver. After infection, mice were fed normal chow or HFD. The weight and fat mass of the mice were measured weekly, and the mice were sacrificed 12 weeks after virus infection (Fig 2A). Overexpression of BATF increased hepatic BATF protein expression (Fig 2B, C), but there was no change in heart, muscle, fat or spleen (Fig 2D, E). In addition, In the case of no difference in feed intake among the four groups (Fig S2A), HFD increased body weight and adiposity in mice, while BATF significantly alleviated weight gain and fat accumulation (Fig 2F, G). Mouse livers showed that HFD made them yellow and covered with fat-like particles, but BATF significantly alleviated these changes (Fig 2H). Liver hematoxylin and eosin (HE) staining of mice in the HFD group confirmed that BATF reduced the number of unstained vacuoles (lipid droplets) (Fig 2I). To verify the effect of BATF on hepatic lipid metabolism, oil red O staining, TG content and Glycerin level were examined and confirmed that BATF significantly alleviated HFD-induced lipid deposition (Fig 2J–M). BATF had no effect on the liver total cholesterol (TC) content of mice (Fig S2B). BATF did not have a significant impact on glucose metabolism in mice, including blood glucose levels (Fig S2C), Glucose Oxidase activity (Fig S2D) and glucose tolerance (Fig S2E, F).

**Figure 2.**
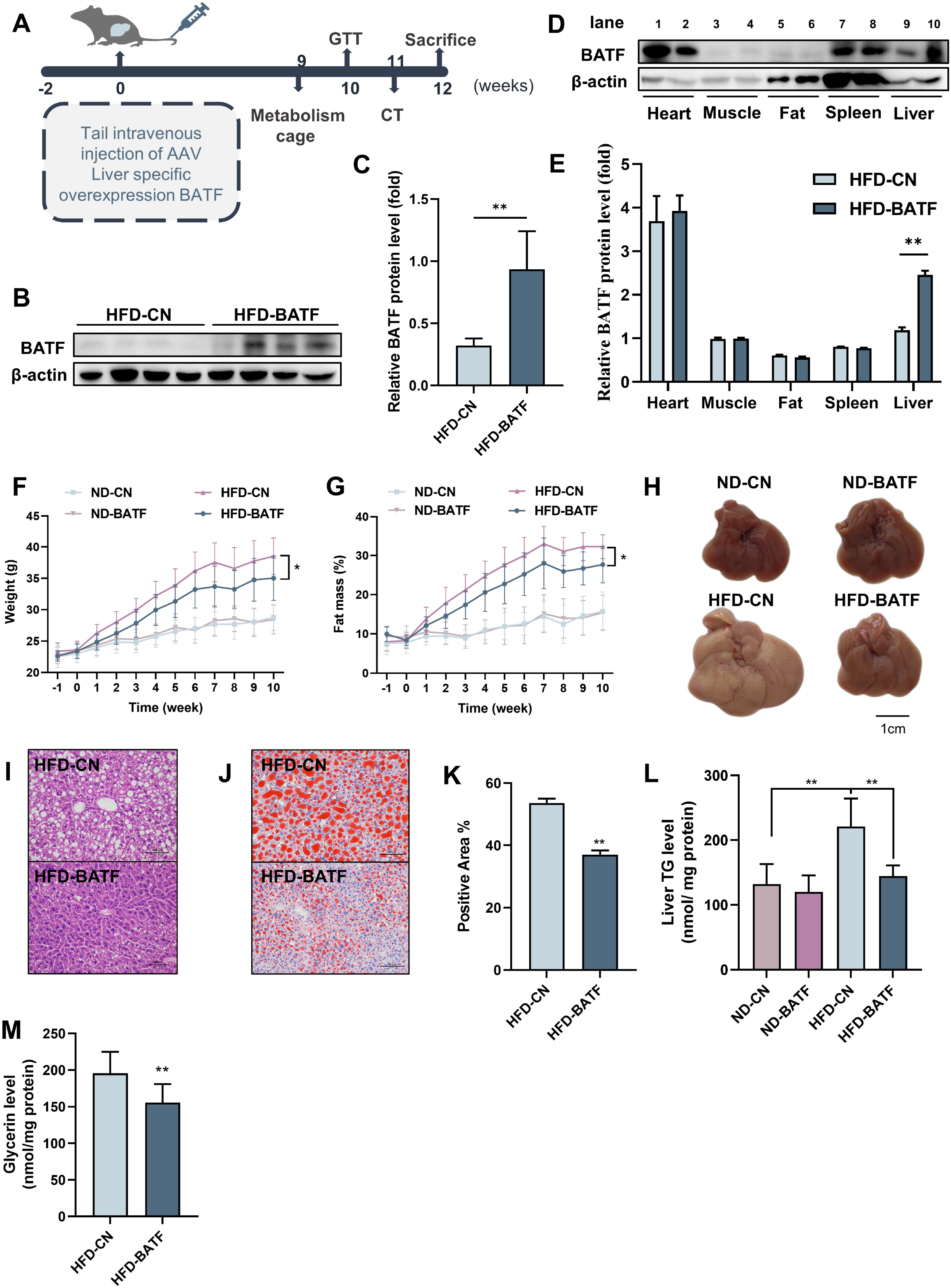
Effects of BATF on hepatic fat deposition in mice. (A) Experimental designs illustration of mice. (B) Expression of BATF protein in liver, (n = 4). (C) Densitometric quantification of the blotting. (D) Expression of BATF protein in various tissues of HFD-CN mice (HBAAV/8-ZsGreen, WB in lane1, 3, 5, 7, 9) and HFD-BATF mice (HBAAV2/8-CMV-m-BATF-3×flag-ZsGreen, WB in lane 2, 4, 6, 8, 10). (E) Densitometric quantification of the blotting. (F) Mice bodyweight, (n = 8-10). (G) Mice fat ratio, (n = 8-10). (H) Mice liver. Bar, 1 cm. (I) HE staining of mice liver sections. Bar, 100 μm. (J) (K) Oil red O staining of mice liver sections and quantitative analysis. Bar, 100 μm, (n = 3). (L) Liver triglyceride levels, (n = 8-10). (M) Liver total Glycerin levels, (n = 8-10).

We conducted further studies to explore the mechanism by which BATF regulated hepatic lipid metabolism. BATF reduced alanine aminotransferase (ALT) and aspartate aminotransferase (AST) activities (Fig 3A, B). It confirmed that BATF alleviated HFD-induced liver injury in mice. Detection of mRNA expression of lipid metabolism genes revealed that BATF did not affect lipid-synthesis-related genes (Fig 3C), but promoted expression of various lipid hydrolysis genes, including *AMPK*α*1, Aco*, *Acox1*, *Bcl2*, *Cpt1*, etc (Fig 3D). Fatty acid β-oxidation is a main route of fat metabolism, in which short-chain acyl coenzyme A dehydrogenase (SCAD) plays a key role. The result confirmed that BATF greatly promotes SCAD activity (Fig 3E). Fatty acid oxidation mainly takes place in mitochondria, so studies on the effect of BATF on aerobic oxidation were carried out. BATF elevated the level of ATP (Fig 3F). An oxygen consumption rate test was performed to explore the effect of BATF on mitochondrial oxygen consumption rate (Fig 3G), and indicated that BATF increased basal respiration, maximal respiration and ATP production in HepG2 cells (Fig 3H). These results suggest that BATF mitigates the effects of HFD on hepatic lipid deposition by promoting energy metabolism.

**Figure 3.**
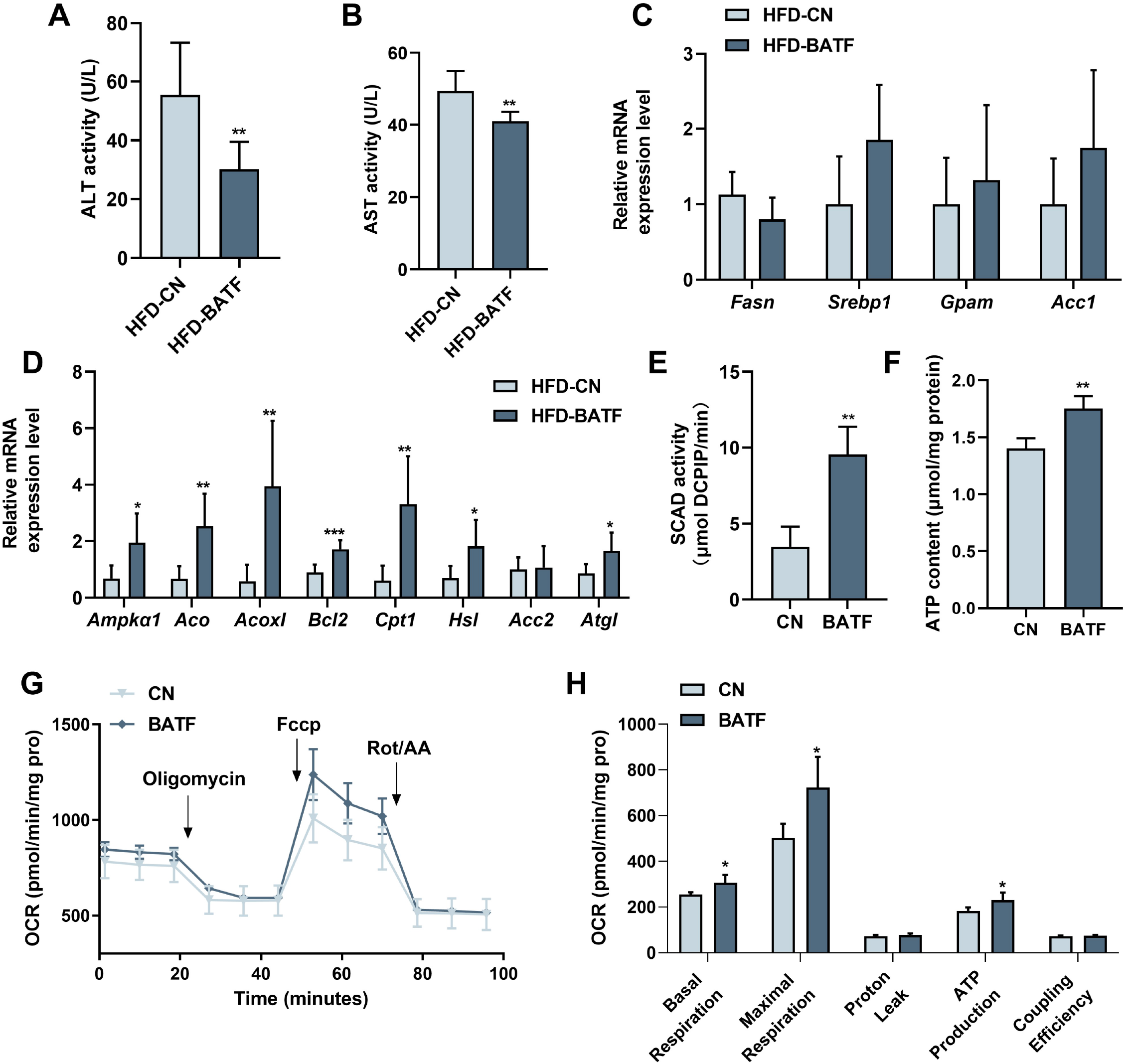
(A) ALT activity in mice liver, (n = 8). (B) AST activity in mice liver, (n = 7). (C) The *Fasn*, *Srebp1*, *Gpam*, *Acc1* mRNA expression level in mice liver, (n = 6-8). (D) The *AMPK*α*1*, *Aco*, *Acox1*, *Bcl2*, *Cpt1*, *Hsl*, *Acc2*, *Atgl* mRNA expression level in mice liver, (n= 6-7). (E) SCAD activity in HepG2 cells with OA/PA treatment, (n = 4). (F) ATP content in HepG2 cells with OA/PA treatment, (n = 4). (G) Oxygen consumption rate (OCR). (H) Basal respiration, maximal respiration, proton leak and coupling efficiency. The data are expressed as mean ± SD. *P < 0.05 **P < 0.01.

### BATF alleviates HFD-induced adipocyte hypertrophy in mice

BATF alleviated HFD-induced increase in adiposity (Fig 2G). After anesthetizing the mice, we used micro-computed tomography (CT) to scan the whole body and image the adipose tissue (Fig 4A). The red lightly shaded parts represent the adipose tissue of the mice, and the results indicated that BATF alleviated overall fat accumulation. The epididymal fat and subcutaneous fat of mice were separated. The results showed that BATF reduced the epididymal and subcutaneous fat (Fig 4B–E). HE staining of mouse adipose tissue allowed microscopic observation of adipocytes, and the diameter and area of the adipocytes indicated that BATF alleviated adipocyte hypertrophy (Fig 4F–K). BATF was overexpressed only in the liver but not in adipose tissue. To investigate the effect of BATF on lipid accumulation in adipocytes, we overexpressed BATF in 3T3-L1 cells and found that it had no effect on accumulation of TG (Fig 4L). These results confirm that BATF does not directly act upon adipocytes to reduce fat accumulation. Therefore, we speculated that BATF regulates fat accumulation in adipose tissue by influencing secretion of substances from the liver. To test this conjecture, we cultured differentiated 3T3-L1 cells with HepG2- cultured cell culture medium (overexpressing BATF) and assayed their TG content. The HepG2-cultured cell culture medium (overexpressing BATF) alleviated TG accumulation in 3T3-L1 cells (Fig 4M). Interleukin (IL)-27 has an inhibitory effect on fat accumulation. We confirmed that HFD decreased IL-27 expression, while BATF enhanced IL-27 expression (Fig 4N). This suggests that BATF mitigates the expansion of adipose tissue by promoting IL-27 secretion.

**Figure 4.**
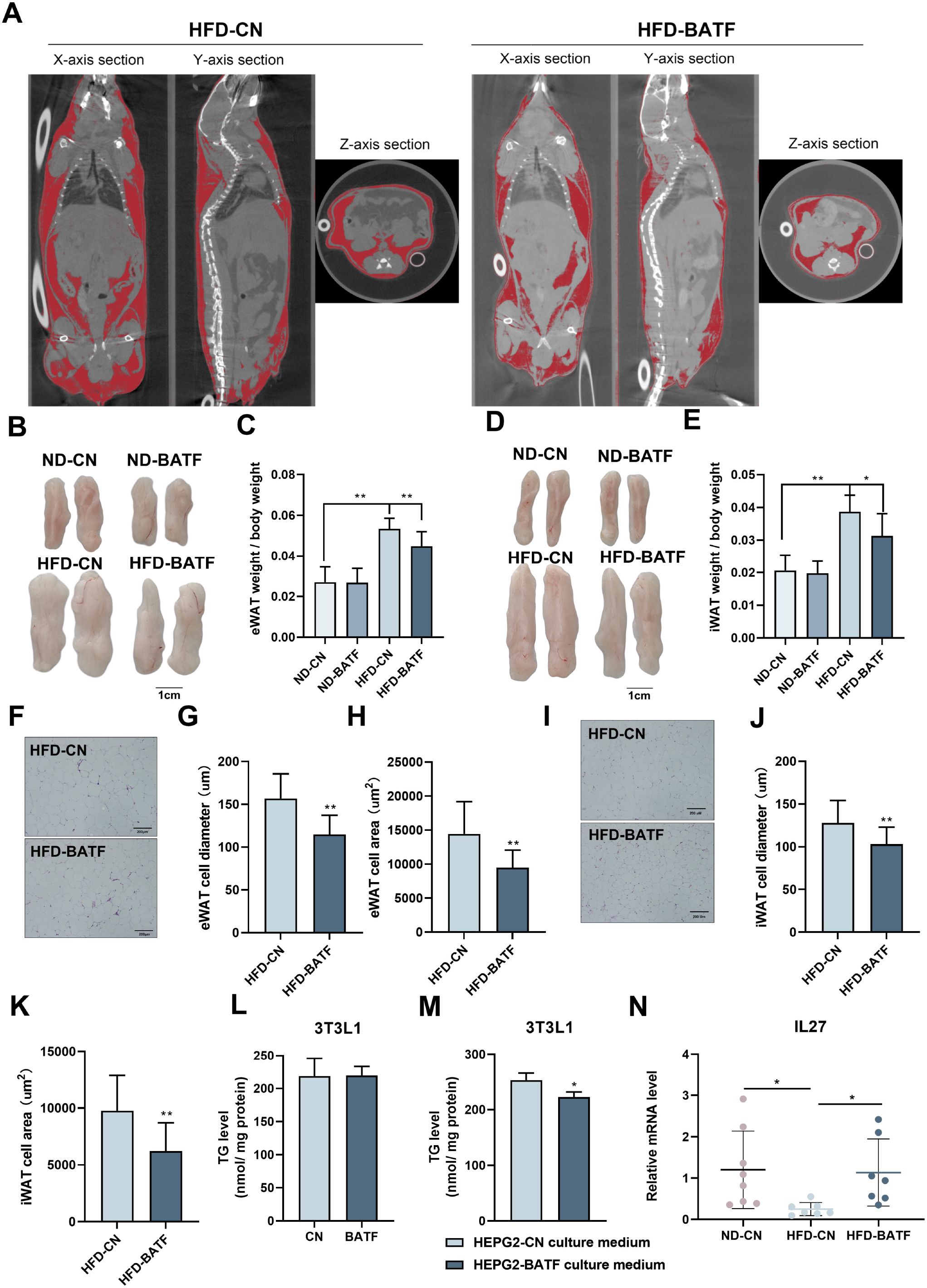
(A) CT images of fat axial view. (B) eWAT of mice. Bar, 1 cm. (C) eWAT weight / bodyweight, (n = 9-10). (D) iWAT of mice. Bar, 1 cm. (E) eWAT weight / bodyweight, (n = 9-10). (F) HE staining of eWAT, (G) adipocyte diameter and (H) cell area. Bar, 200 μm. (I) HE staining of iWAT, (J) adipocyte diameter and (K) cell area. Bar, 200 μm. (L) Triglyceride content of undifferentiated 3T3L1 cells (n=5). (M) Triglyceride content of differentiated 3T3L1 cells (n=3-4). (N) The mRNA expression of IL27 in liver tissues (n = 5). The data are expressed as mean ± SD. *P < 0.05 **P < 0.01.

### BATF alleviates hepatocyte lipid accumulation by inhibiting programmed cell death protein 1

Several studies have confirmed that there is a regulatory relationship between BATF and programmed cell death protein (PD)1^14–16^. HFD increased PD1, while BATF inhibited PD1 expression in mouse liver (Fig 5A). These results were confirmed in HepG2 cells (Fig 5B). To determine the role of PD1 in lipid accumulation in hepatocytes, we overexpressed PD1 in HepG2 cells and assayed the lipid content. Both oil red O staining and TG content confirmed that PD1 promoted lipid deposition in hepatocytes (Fig 5C, D). PD1 antibody inhibited the role of PD1 and successfully alleviated TG accumulation in hepatocytes, but BATF overexpression plus PD1 antibody did not further alleviate lipid deposition (Fig 5E), suggesting that BATF regulates lipid deposition through PD1. To confirm this hypothesis, we overexpressed BATF and PD1. Overexpression of BATF decreased TG, while the TG-lowering effect of BATF disappeared when PD1 was overexpressed (Fig 5F). To determine the regulatory role of BATF on PD1, we constructed a luciferase vector containing 2000 bp upstream of the transcription start site of PD1. BATF decreased the transcriptional activity of PD1 promoter, confirming the role of BATF in regulating PD1 expression (Fig 5G). These results suggest that BATF regulates lipid deposition through inhibition of PD1. In conclusion, HFD leads to an increase in the level of PD1 transcription in the liver. BATF promotes lipolysis and energy consumption in hepatocytes by regulating PD1 transcription, thereby reducing HFD-induced liver lipid deposition and alleviating NAFLD (Fig 5H).

**Figure 5.**
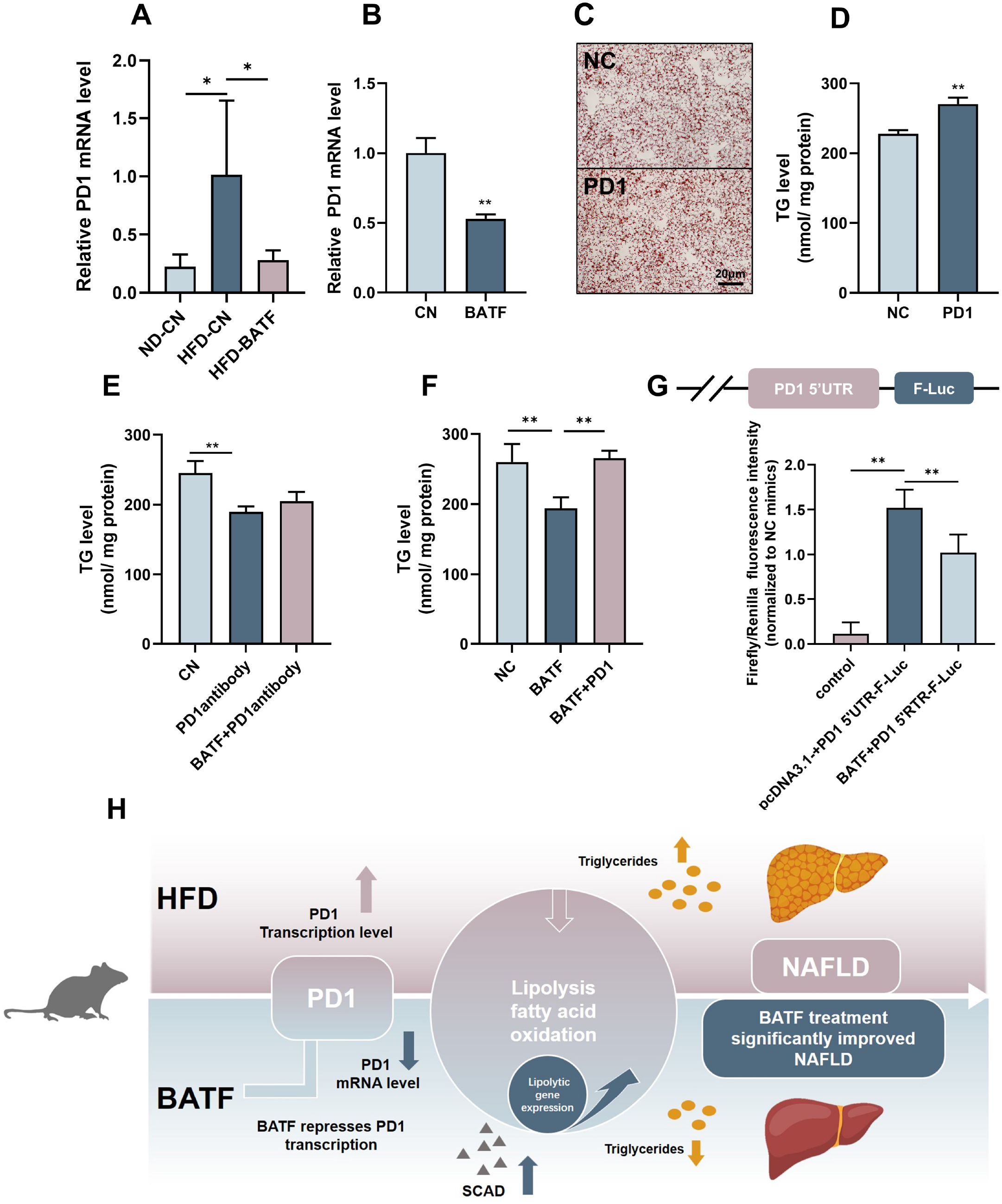
BATF alleviates hepatocyte lipid accumulation by inhibiting PD1. (A) The mRNA expression of PD1 in liver tissues (n = 3). (B) The mRNA expression of PD1 in HepG2 cells (n = 3). (C) Oil red O staining of HepG2 cells, Bar, 20 μm, (n = 3). (D), (E), (F) Triglyceride content with OA/PA (n ≥ 3). (G) Dual luciferase assay on Hepa1-6 cells cotransfected with firefly luciferase constructs containing the PD1 promoter, Renilla luciferase vector pRL-TK and pcDNA3.1(-) or pcDNA3.1(-)-BATF, (n ≥ 3). (H) The Mechanism diagram of BATF alleviates hepatocyte lipid accumulation by PD1. The data are expressed as mean ± SD. *P < 0.05 **P < 0.01.

### PD1 antibody alleviates HFD-induced obesity and liver steatosis in mice

PD1 plays an important role in tumorigenesis as a tumor immune escape target, and PD1 antibodies as antitumor drugs can inhibit PD1 and mitigate tumor growth and cancer development^17^. Our study confirmed that PD1 promoted lipid deposition in hepatocytes (Fig 5C, D), and *in vivo* experiments were performed to verify the role of PD1 on hepatic steatosis in mice fed an HFD. PD1 antibody alleviated HFD-induced weight gain in mice after intraperitoneal injection of PD1 antibody (200 μg) (Fig 6A). The mice injected with PD1 antibody were smaller than the control mice (Fig 6B). The results of fat mass confirmed that PD1 antibody alleviated fat accumulation in mice (Fig 6C). The epididymal and subcutaneous fat of mice were reduced by PD1 antibody (Fig 6D–G). Mouse livers showed that HFD gave them an appearance of hepatic steatosis, which was significantly alleviated by PD1 antibody (Fig 6H). The liver was stained with Oil Red O and assayed for TG content, which showed that PD1 antibody reduced accumulation of lipid droplets (Fig 6I) in the liver and alleviated lipid deposition (Fig 6J). We extracted RNA from mouse livers and examined expression of lipolysis-related genes. PD1 antibody treatment was consistent with overexpression of BATF, and both promoted expression of lipolysis-related genes (Fig 6K). Similarly, PD1 antibody treatment promoted SCAD activity in mouse liver, suggesting fatty acid β-oxidation was enhanced (Fig 6L).

**Figure 6.**
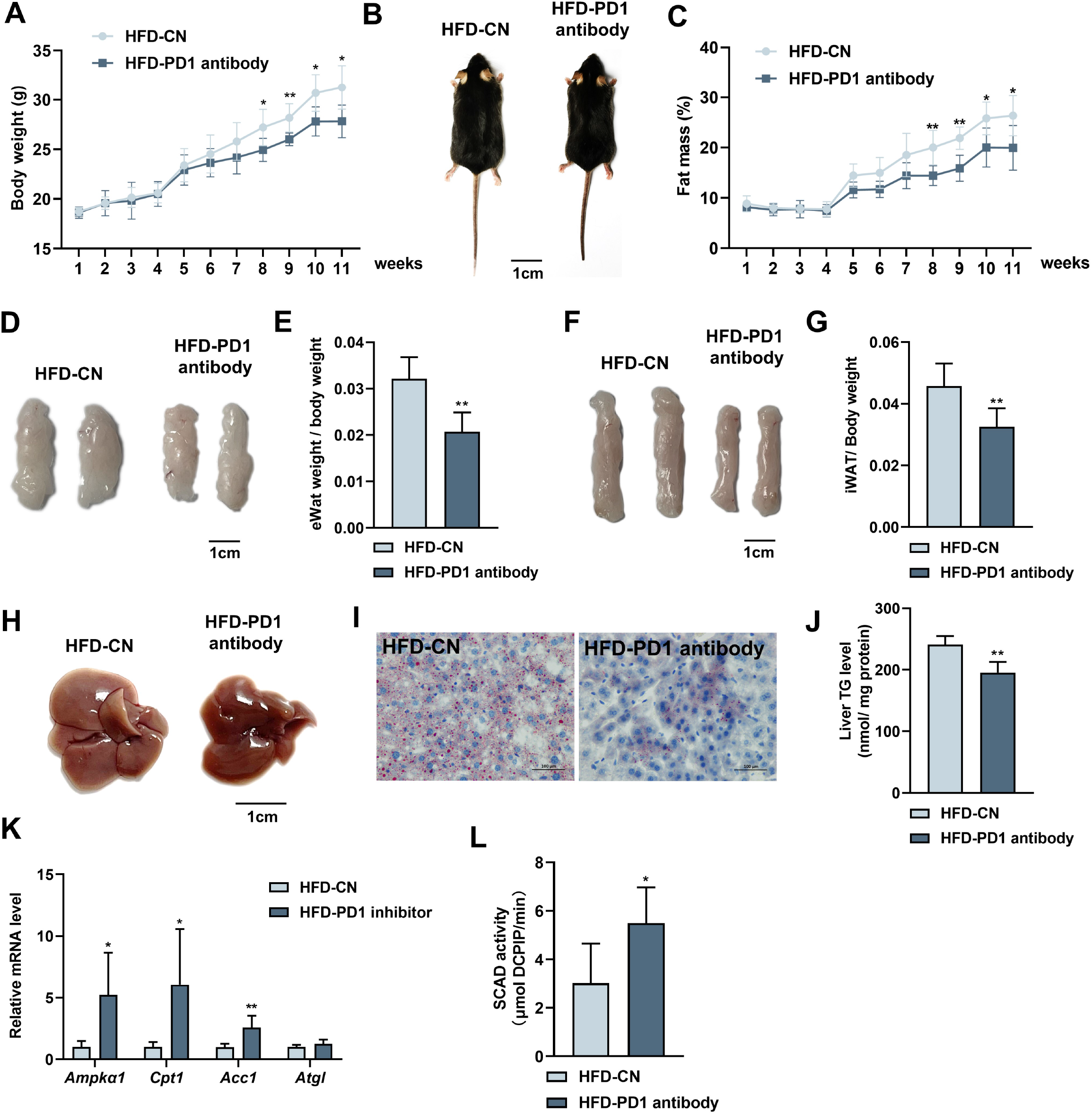
Effects of PD1 antibody on liver lipid metabolism in HFD mice. (A) Mice bodyweight, (n ≥ 5). (B) The mice were injected with IgG or PD1 antibody under HFD, (n ≥ 5). (C) Mice fat ratio, (n ≥ 5). (D) eWAT of mice. Bar, 1 cm. (E) eWAT weight / bodyweight, (n ≥ 5). (F) iWAT of mice. Bar, 1cm. (G) eWAT weight / bodyweight, (n ≥ 5). (H) The liver of mice. (I) HE staining of mice liver sections. Bar, 100 μm. (J) Liver triglyceride (TG) levels. (K) The *AMPK*α*1*, *Cpt1*, *Acca*, *Atgl* mRNA expression level in mice liver, (n ≥ 5). (L) SCAD activity in liver of mice, (n = 5). The data are expressed as mean ± SD. *P < 0.05 **P < 0.01 .

## Discussion

In the present study, we found that HFD elevated BATF expression in mouse liver. We also found that inhibition of hepatocyte BATF did not regulate lipid deposition, while overexpression of BATF decreased TG accumulation. Studies on hepatocytes confirmed that BATF regulated hepatocyte lipid deposition. BATF also alleviated HFD-induced obesity in mice, not by directly acting on adipose tissue but by influencing IL-27, which acts as a regulator of fat deposition^18^. The regulation of hepatic lipid metabolism by BATF has not previously been investigated. Here, we found that BATF mitigated HFD-induced increase in PD1 expression. We revealed that PD1 promoted hepatocyte lipid deposition. Through luciferase assays, we found that BATF regulated lipid deposition by inhibiting PD1 expression. Experiments in mice confirmed that PD1 inhibition alleviated HFD-induced obesity and hepatic steatosis.

The adaptive immune system has evolved to eliminate virtually any threat from the organism^19^. PD1 is expressed during T-cell activation and through T-cell receptor counteracting positive signaling by engaging its ligands PDL1 and/or PDL2^20^. PD1 regulates the immune system and promotes self-tolerance by downregulating the immune response to human cells and by suppressing T-cell inflammatory activity^21^. The development of checkpoint blockade inhibitors has revolutionized cancer immunotherapy^22^ and has led to durable survival outcomes in some patients with metastatic disease^23^. Previous studies on PD1 have focused on the immune system, and we found that HFD increased its expression in the liver of mice (Fig 5A). This is consistent with previous studies^24^ suggesting that PD1 is equally important in the liver. Several studies have demonstrated that PD1 and BATF regulate each other to modulate immune system flexibility^25–26^. We hypothesized that BATF and PD1 co-regulate the hepatic immune system to affect lipid deposition. Our results confirm our conjecture that BATF reduces fat accumulation by inhibiting PD1, confirming that PD1 is downstream of the BATF signaling pathway (Fig 5F, G).

Cancer and autoimmune disease are closely related, and many therapeutic antibodies are widely used in clinics for the treatment of both diseases. Moreover, immune checkpoint blockade using the anti-PD1/PD-L1/CTLA4 antibody has been shown to improve prognosis of patients with refractory solid tumors^27^. In this study, we used PD1 antibodies to treat HFD-induced obesity and NAFLD in mice. Surprisingly, PD1 antibody has an excellent therapeutic effect on obesity and NAFLD. PD1 antibodies reduced adiposity and hepatic lipid deposition (Fig 6), which is consistent with overexpression of BATF. There have been no approved drugs for NAFLD treatment. In this study, the effect of PD1 antibodies on NAFLD treatment was encouraging. Our findings have somewhat pioneered the cognition of BATF, PD1 and its expansion with NAFLD therapy.

## Materials and methods

### Animals

All animal studies were approved by the Animal Ethics Committee of Guangxi University (GXU-2020-288). Four-week-old male C57BL/6J mice were purchased from Guangxi Medical University (Nanning, China) and kept in individual cages on a 12-h light/dark cycle with free access to water and food at room temperature. At week 10, mice were injected intravenously (tail vein) with 200 μl adeno-associated virus (AAV). AAV was HBAAV2/8-CMV-m-BATF-3×flag-ZsGreen and HBAAV/8-ZsGreen (packaged by HANBIO, Shanghai, China). The mice were fed with a normal diet (ND) (AIN93) or HFD (D12492) for 12 weeks^28^. To study the effect of PD1 inhibitors on NALFD, the mice were fed with HFD and rat IgG (ab37361; Abcam) or anti-mouse PD1 (AB10949053, Bio X Cell) antibody was administered (intraperitoneal injection) at 200 μg/mouse twice weekly^29^. Body weight and fat mass were recorded weekly, and body composition (fat) was determined by Body Composition Analyzer (QMR-23-060H-I; Suzhou, China). Mice were anesthetized with 2.4% tribromoethanol (T48402, Sigma–Aldrich) to evaluate fat content, and micro-CT images were taken using a micro-CT scanner (SKYSCAN1278; Bruker, Belgium). All groups were randomized and blinded to form equal-sized groups of at least three (exact numbers are given in the legend).

### Human disease data

The human data of NAFLD were taken from the Gene Expression Omnibus, Accession Number GSE130970^30^.

### Cell culture and transfection

HepG2 (HB-8065, ATCC), L02 (YUCHI Biology, China), Hepa1-6 (CRL-1830, ATCC), HEK293T (ACS-4500, ATCC), 3T3L1 (CL-173, ATCC) cells were cultured in DMEM medium (C11995500BT, Gibco) supplemented with fetal bovine serum (04-400-1A, BI) and 1% penicillin–streptomycin (P1400, Solarbio, China). The AML12 cell (CRL-2254, ATCC) culture medium was DMEM/F12 (Gibco) containing 10% FBS and 1% penicillin–streptomycin. The cells were incubated at 37°C and 5% carbon dioxide. For simulating HFD *in vitro*, 200 μM OA and 100 μM PA were conjugated to bovine serum albumin (BSA) as previously described^31^.

For transfection of plasmids or siRNA oligos, Hieff Trans Liposomal Transfection Reagent (40802ES02, YEASEN) was used. Plasmid constructs were created using the eukaryotic expression vector pcDNA3.1-^32^. Mice BATF (NM_016767.2), PD1 (NM_008798.3) CDS were cloned from mouse liver cDNA. RNAi sequence was synthesized from Sangon Biotech (Shanghai, China).

### Primary hepatocyte isolation and AAV infection

The abdominal cavity of 8-week-old male mice was opened after anesthesia with 2.4% tribromoethanol. The inferior vena cava was cannulated with a 23G needle and perfused with 30 mL perfusion buffer (1×HBSS without Ca^2+^, Mg^2+^ supplemented with 0.5 mM EDTA and 25 nM HEPES). The liver was digested by 25 mL of 37 [digestion buffer (1×HBSS with Ca^2+^, Mg^2+^ supplemented with 1 μg/mL collagenase type II and 1M HEPES). Following the perfusion, the liver was gently removed and massaged through 70 μm nylon mesh with 10 mL of 4 [precooled complete medium (DMEM supplemented with 1% FBS). Hepatocytes were isolated by centrifugation at 50 *g*, 4°C, for 2 min. After discarding the supernatant, hepatocytes were resuspended in 10 mL complete medium containing 5 mL freshly prepared 90% Percoll solution (9 mL Percoll with 1 mL PBS). Cell debris was removed by centrifugation at 200 *g*, 4°C, for 10 min. The supernatant was aspirated, resuspended and centrifuged at 50 *g*, 4 [for 2 min, after resuspension in William E medium supplemented 1% glutamine and 1% penicillin–streptomycin. Cell resuspension was transferred into a 0.1% gelatin-coated plate and incubated at 37°C, 5% carbon dioxide^33^. Cells were maintained for 24 h in medium. For *in vitro* experiments with AAV, 1 μL f purified HBAAV2/8-CMV-m-BATF-3×flag-ZsGreen or HBAAV/8-ZsGreen was added to cultured hepatocytes to achieve high BATF overexpression.

### Glucose tolerance test

Glucose tolerance test (GTT) was performed as previously described^34^. Mice were fasted for 16 h and injected with D-glucose at 2 g/kg intraperitoneally. The blood glucose level was measured at 0,15, 30, 60 and 120 min after glucose injection.

### Western blotting

Tissues were lysed with 1 mL RIPA lysis buffer and supplemented 1% protease inhibitor cocktail (Solarbio) for 2–5 min with steel balls in a tissue lyser (Qiagen, Germany). Tissue lysates were incubated with the buffer at 4°C overnight (cell samples were incubated in lysis buffer for 30 min at 4°C). The lysate was centrifuged at 12,000 rpm for 10 min, the supernatant was collected, and the protein concentration was determined using the BCA method (P0011, Beyotime Biotechnology). The supernatant was mixed with loading buffer and boiled at 100°C for 10 min. Samples were loaded and subjected to 10% SDS-PAGE. After transfer to PVDF membranes, blots were blocked with 5% skimmed milk powder and incubated with primary antibody overnight at 4°C. The primary antibodies included a BATF antibody (8638S, Cell Signaling Technology, USA), anti-β-actin (GB11001, Servicebio, China), and β-tublin (15115, Cell Signaling Technology). Secondary antibody was goat anti-rabbit (Jackson, 111-035-003), images were acquired using the BIO-RAD protein gel imaging system (Universal hood II), and band intensity was calculated using Image J^35^.

### TGs, TC and glycerin detection

Cells or tissues were prepared according to the instructions of the A110-1-1 Triglyceride Assay Kit^36^, A111-1-1 Total Cholesterol Assay Kit (Nanjing Jiancheng Bioengineering Institute, China), and Tissue Cell Glycerin Test kit (E1024, Pulilai, China). Liver tissues or cells were lysed by RIPA lysis buffer. Samples were centrifuged at 12 000 rpm for 10 min, and the supernatants were collected. Supernatants (10 μL) were added to a 96-well plate and 190 μL working solution was added to each well. The plate was incubated at 37°C for 10 min. The microplate reader (Tecan M200 PRO) recorded the contents of TG, TC and glycerin, then normalized using the protein concentration.

### Enzyme activity assay

SCAD enzyme activity in cultured cells or liver tissues was detected using the SCAD Assay Kit (Gen Med, Plymouth, MN, USA). The liver tissue activities of ALT and AST were measured using the C009-2-1 ALT Reagent Kit and C010-2-1 AST Reagent Kit (Nanjing Jiancheng Bioengineering Institute, China). The liver tissue activities of glucose oxidase was measured using the BC0695 Glucose Oxidase(GOD) Activity Assay Kit (Servicebio, China).

### ATP measurements

Cellular ATP content was measured using a luciferin/luciferase-based kit (S0026, Beyotime Biotechnology, China).

### HE and Oil Red O staining

HE and Oil Red O staining was performed as previously described^37^. Livers, epididymal fat and subcutaneous fat of mice were dissected and fixed in 4% paraformaldehyde overnight at 4°C. Samples were sent to Wuhan Service Technology Co. Ltd. (Wuhan, China) for paraffin embedding, sectioning, and staining. Images were collected by light microscope and Image J was used to examine the cell diameter and the area.

### Oxygen consumption rate

Oxygen consumption rate was measured by extracellular flux (XF24) analyzer (Seahorse Bioscience). Cells were seeded at 2 × 10^4^ per well in XF24 plates. The probe plate was hydrated with the calibration solution (pH 7.4) and a 2 mM glutamine assay solution (pH 7.35 ± 0.05) was prepared, and incubated for 16–20 h at 37°C and 5% CO_2_. Culture medium was replaced by 2% FBS medium and incubated without CO_2_ for 1 h before the completion of probe cartridge calibration. Basal oxygen consumption rate was measured after injection of oligomycin (1 μM), carbonyl cyanide p-trifluoro-methoxyphenyl hydrazone (FCCP 1 μM), and rotenone/antimycin A (ROT/AA) (0.5 μM)^38^.

### RNA extraction and real-time quantitative polymerase chain reaction

Total RNA was extracted by TRIzol Reagent (Life Technologies, USA). Reverse transcription of RNA was performed with the Reverse Transcription System (A3500, Promega), and formed cDNA from 1 μg total RNA. 2× RealStar Green Fast Mixture was used for real-time quantitative polymerase chain reaction (PCR) (GenStar, Beijing, China). The relevant primer sequences (5′-3′) were as follows: Human β-actin forward: CACCATTGGCAATGAGCGGTTC, reverse: AGGTCTTTGCGGATGTCCACGT. Human BATF forward: GATGTGAGAAGAGTTCAGAGGAG, reverse: GTTTCTCCAGGTCTTCGCTCTC. Mouse BATF forward: ATGCCTCACAGCTCCGACAGC, reverse: TCAGGGCTGGAAGCGTGGC. Mouse PD-1 forward: CGGTTTCAAGGCATGGTCATTGG, reverse: TCAGAGTGTCGTCCTTGCTTCC; Mouse *Fasn* forward: CACAGTGCTCAAAGGACATGCC, reverse: CACCAGGTGTAGTGCCTTCCTC. Mouse *Srebp1* forward: CGACTACATCCGCTTCTTGCAG, reverse: CCTCCATAGACACATCTGTGCC. Mouse *Gpam* forward: GCAAGCACTGTTACCAGCGATC, reverse: TGCAATCAGCCTTCGTCGGAAG. Mouse *Acc1* forward: GTTCTGTTGGACAACGCCTTCAC, reverse: GGAGTCACAGAAGCAGCCCATT. Mouse AMPKα1 forward: GGTGTACGGAAGGCAAAATGGC, reverse: CAGGATTCTTCCTTCGTACACGC. Mouse Aco forward: TCACAGCAGTGGGATTCCAA, reverse: TCTGCAGCATCATAACAGTGTTCTC. Mouse *AcoxI*forward: TAACTTCCTCACTCGAAGCCA, reverse: AGTTCCATGACCCATCTCTGTC. Mouse *Bcl2* forward: ACTTCCACTACAGGACAGAC, reverse: TCTAAGGTGACTCGATATGG. Mouse *Cpt1* forward: GCACTGCAGCTCGCACATTACAA, reverse: CGTTGACATCCGTAAAGACC. Mouse β*-actin* forward: AACAGTCCGCCTAGAAGCAC, reverse: CTCAGACAGTACCTCCTTCAGGAAA. Mouse *Hsl* forward: GGAGCACTACAAACGCAACGA, reverse: TCGGCCACCGGTAAAGAG. Mouse Acc2 forward: AGAAGCGAGCACTGCAAGGTTG, reverse: GGAAGATGGACTCCACCTGGTT. Mouse Atgl forward: GGAACCAAAGGACCTGATGACC, reverse: ACATCAGGCAGCCACTCCAACA.

### Retroviral transduction

Retroviral vectors pMSCV-PIG and pMSCV-PIG-BATF were used to infect 3T3L1 cells. The pMSCV-PIG-BATF viral vector was constructed, and the coding sequence of mouse BATF was amplified from liver cDNA by PCR and cloned into the *Eco*RI site using ClonExpress II One Step Cloning Kit (C112-01/02, Vazyme, China). To obtain pseudotyped virus with the ability to infect, pMSCV-PIG-BATF, pMSCV-PIG-GFP, and retroviral packaging plasmids (pKat and pCMV-VSV-G) were cotransfected into subconfluent HEK293T cells. The virus stocks were collected 2 and 3 days after transfection, centrifuged at 5000 *g* for 10 min, subpackaged and frozen at −80°C. 3T3L1 cells with around 30% confluence were treated with virus stock containing 8 μg/mL polybrene for 4 h, washed, and cultured in fresh medium.

### Luciferase assay for promoter activity analysis

Firefly luciferase constructs were cloned using the pGL3-Basic vector as a backbone. Target 5′ UTRs (PD1 was amplified by primers, forward: AGCATGAGCCCTGAGGATTG reverse: CAGTGTCGCCTTCAGTAGCA) were cloned upstream of Firefly luciferase in the pgl3. Renilla luciferase reporter vector pRL-TK was gifted by State Key Laboratory for Conservation and Utilization of Subtropical Agro-Bioresources. Hepa1-6 cells were inoculated into medium in 24-well culture plates and cultured overnight. Cell transfection and cotransfection were conducted when the cell density reached 80%. Dual Luciferase Reporter Assay was performed 48 h post-transfection using the Dual-Lumi II Luciferase Reporter gene Assay kit (PG089S). Renilla luciferase was used to normalize transfection efficiency and luciferase activity.

### Statistical analysis

All experiments were repeated at least in triplicate, and statistical analysis was done using independent values. All data are presented as mean[±[SD. Statistical analysis was performed using Student’s unpaired two-tailed *t* test (for two groups) or analysis of variance (ANOVA) (for multiple groups). One-way ANOVA with Tukey’s test was conducted for comparisons. The differences were considered significant at P[<[0.05 (*). Values with different letters were significantly different.

## Acknowledgements

This work was supported by National Natural Science Foundation of China (32272952); Guangxi Science Foundation for Distinguished Young Scholars (2020GXNSFFA297008); Guangxi Natural Science Foundation (2019GXNSFDA245029); Guangxi Academy of Medical Sciences high-level Talents Foundation (YKY-GCRC-202302).

---

**Figure S1.**
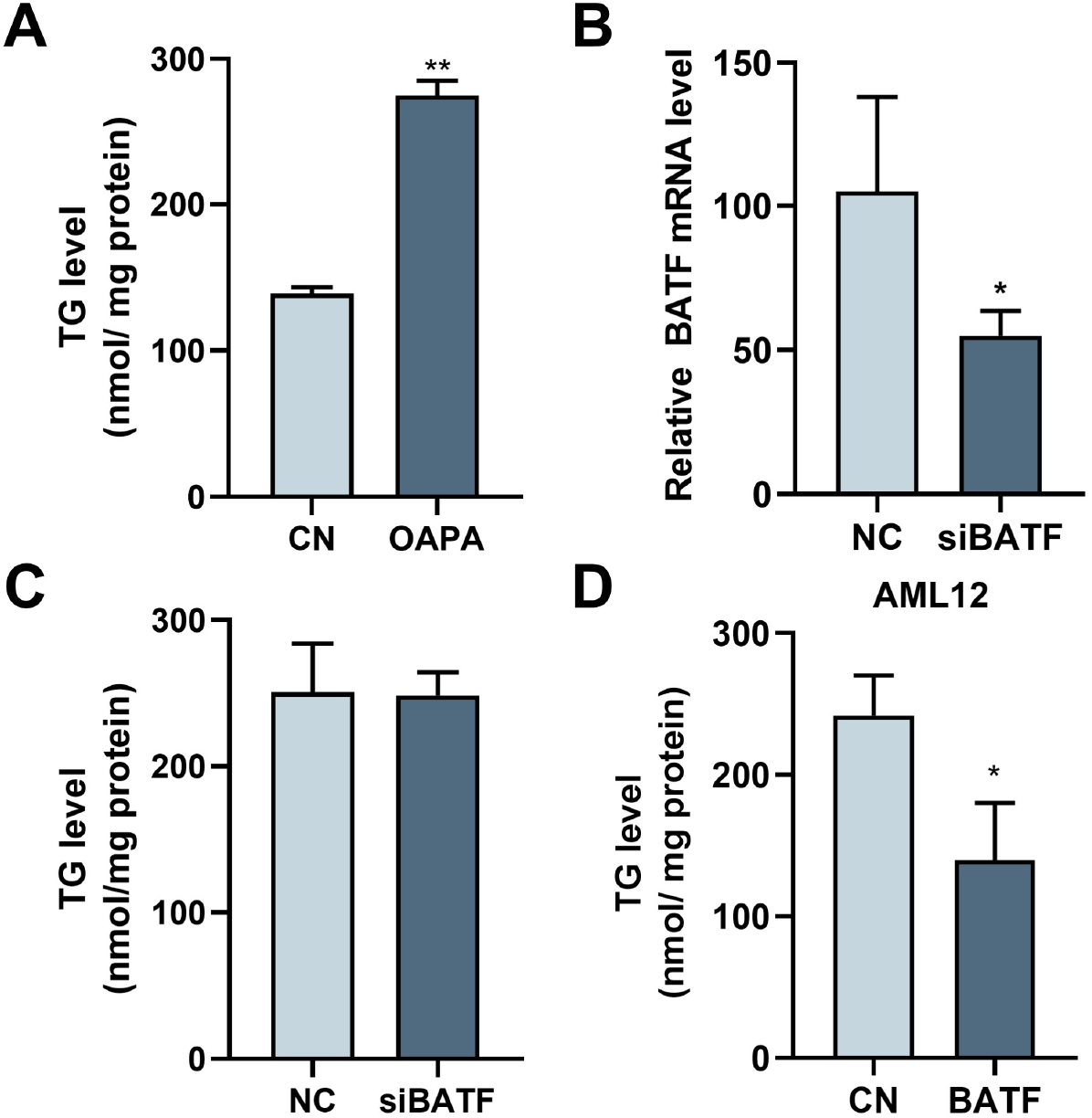
(A) Triglyceride content with OA/PA (n = 3). (B) BATF mRNA levels. NC, negative control group; siBATF, BATF inhibition group, (n = 3). (C) Triglyceride content with OA/PA treatment, (n = 5). (D) Triglyceride content with OA/PA when BATF was overexpressed, (n = 3). The data are expressed as mean ± SD. *P < 0.05 **P < 0.01.

**Figure S2.**
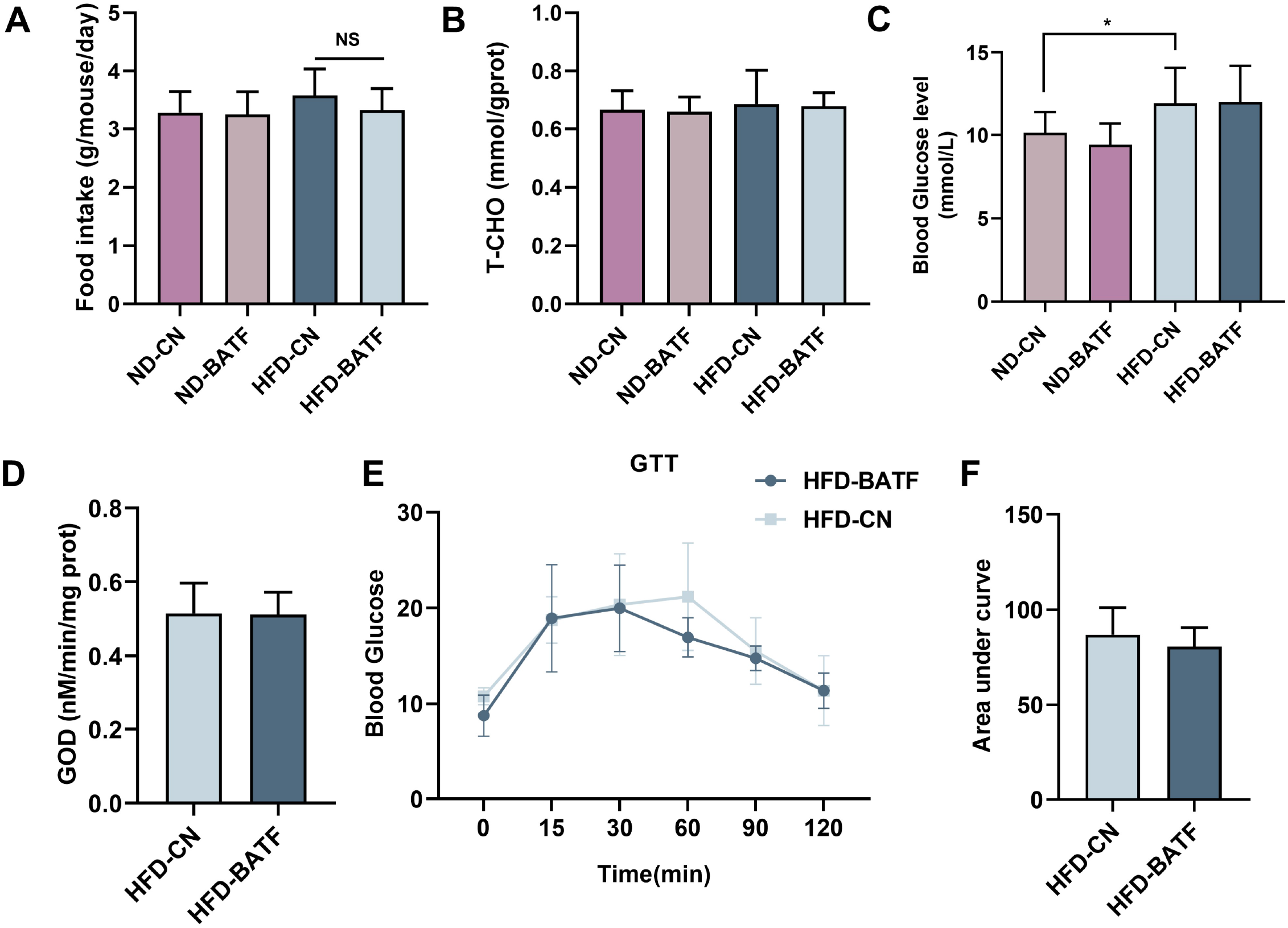
(A) Average daily feed intake (n= 7). (B)Liver total cholesterol levels, (n = 8-10). (C) Fasting blood glucose level in mice, (n = 8-10). (D)Liver glucose oxidase activity. (E) Glucose tolerance test and (F) quantitative analysis, (n = 5-6). The data are expressed as mean ± SD. *P < 0.05 **P < 0.01.

